# The Golgi vesicle tether p115/USO1 can bind directly to the ER exit site organiser Sec16A

**DOI:** 10.1101/2025.10.16.682774

**Authors:** Igor Yakunin, Alison K Gillingham, Conceição Pereira, David C Gershlick, Sean Munro

## Abstract

Newly-made secretory and membrane proteins exit the endoplasmic reticulum (ER) in COPII vesicles that form at specialised ER exit sites. These exit sites are typically near to the early Golgi compartments that receive these vesicles. A key player in the delivery of vesicles to the early Golgi is p115 (USO1), a homodimer with a folded head domain and a coiled-coil tail that is anchored to Golgi membranes. p115 has been shown to capture vesicles and to bind to SNARE proteins to promote membrane fusion. Here we report that the head domain of human p115 can bind directly to Sec16A, a large scaffolding protein that organises ER sites and promotes COPII vesicle formation. Structural prediction and deletion mapping define the region of interaction to a conserved motif in the unstructured N-terminal region of Sec16A, and mutations in p115 that block motif binding reduce the efficiency of secretion. This interaction could potentially allow a subset of p115 molecules to reach across from the early Golgi to the ER exit sites to contribute to the large-scale organisation of the early secretory pathway.

## INTRODUCTION

The Golgi apparatus is the major sorting hub in the secretory pathway. It receives newly made lipids and proteins from the endoplasmic reticulum (ER) and then after they move through the Golgi, they are sorted to the cell surface or the compartments of the endocytic system (Glick and Luini, 2011; Pantazopoulou and Glick, 2019). Export from the ER is mediated by the COPII coat that forms vesicles and other carriers at ER exit sites (Downes and Zanetti, 2025; Gomez-Navarro and Miller, 2016; Peotter et al., 2019). Soon after detaching from the ER, these carriers fuse to the ER-Golgi intermediate compartment (ERGIC) or the cis Golgi where the COPI coat forms vesicles that capture escaped ER residents for return to the ER (Barlowe and Helenius, 2016). The ER exit sites and ERGIC are often closely opposed and there is growing evidence that they are organised as a functional unit that allows the rapid generation and consumption of the COPII and COPI vesicles that mediate anterograde and retrograde traffic respectively (Downes et al., 2025). In mammalian cells there is a further level of organisation as there are ER exit sites throughout the ER network that fills the cytoplasm, whereas the Golgi apparatus is typically concentrated in the perinuclear region. In the cell periphery the ERGIC generates tubular carriers which move on microtubules toward the Golgi, or the ERGIC compartments move en bloc to the Golgi once they reach a certain size (Stephens et al., 2000). Next to the Golgi in the centre of the cell, there is typically a concentration of ER exit sites and the ERGIC that the COPII vesicles fuse with corresponds to the forming cis Golgi compartment of the Golgi stack which also receives the carriers from the peripheral ERGIC.

Several proteins have been proposed to contribute to the organisation of this ER-ERGIC unit of the secretory pathway in addition to the SNARE proteins that drive the fusion of the vesicles with their destinations (Farhan et al., 2025). These include in the ER, the peripheral scaffolding protein Sec16A which helps recruit COPII coats into defined patches on the ER, and the integral membrane protein TANGO1 which projects into both the ER lumen where it is proposed to select specific cargo and into the cytoplasm where it is required to help organise the exit sites (Hughes and Stephens, 2010; McCaughey et al., 2021). The cytosolic protein TFG can bind to COPII subunits and has been proposed to help in forming the COPII coat and also directing the clustering of COPII-coated carriers once they have budded (Hanna et al., 2017). On the ERGIC and cis Golgi various factors are likely to act very early in the transport process. The TRAPPIII complex activates the Rab1 small GTPase which has been proposed to recruit, amongst other effectors, the GBF1 exchange factor that activates Arf proteins to generate retrograde COPI vesicles (Galindo and Munro, 2023; Monetta et al., 2007). The golgin coiled-coil proteins GM130 and GMAP210 are on the ERGIC and cis Golgi and contribute to efficient trafficking (Roboti et al., 2015; Seemann et al., 2000). One role of GM130 is to bind via its basic N-terminus to the acidic C-terminal region of the protein p115 (USO1), although the functional importance of this interaction is unclear as this part of p115 is not required for it to localise to the Golgi or to maintain Golgi structure and protein secretion (Nelson et al., 1998; Puthenveedu and Linstedt, 2004). P115 was first identified in yeast as a temperature sensitive mutant, *Uso1-1*, that blocked ER to Golgi transport (Nakajima et al., 1991). Its mammalian orthologue was then discovered independently as a factor required in two in vitro assays for vesicular transport in the Golgi (Barroso et al., 1995; Sapperstein et al., 1995). It is localized to the ERGIC and the cis face of the Golgi, and of the various putative structural proteins in this region of the pathway, it is the best conserved in evolution and appears to be essential in all organisms tested to date ranging from mammals through to yeast and protozoa (Kim et al., 2012; Pasquarelli et al., 2024; Seog et al., 1994). It has a head domain made up of armadillo repeats and a C-terminal coiled-coil domain that directs dimerization (Fig. 1A,B) (Sapperstein et al., 1995; Yamakawa et al., 1996).

**Figure 1.**
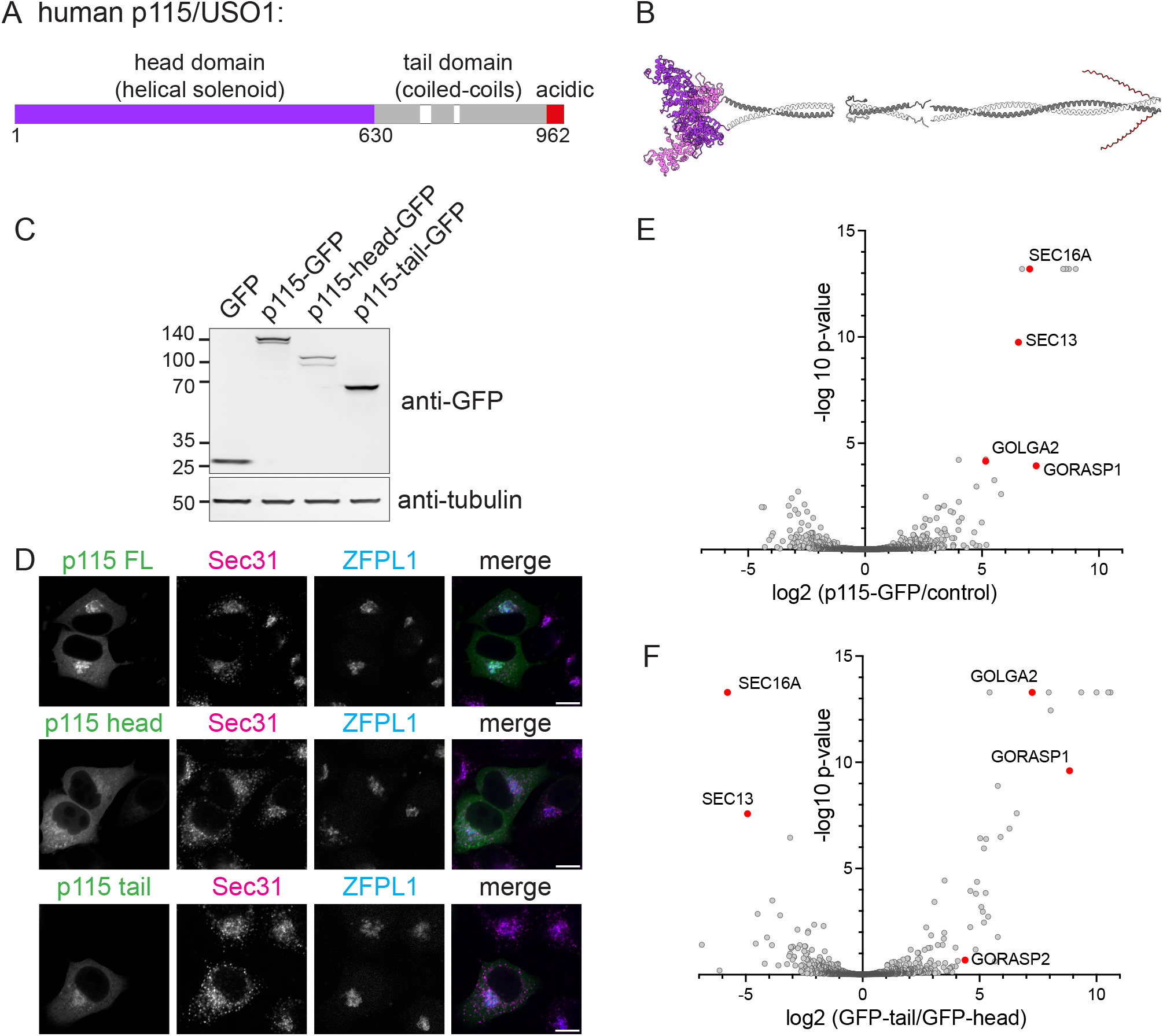
Identification of proteins co-precipitating with p115. **(A)** A schematic of p115: A globular head domain comprised of armadillo repeats is followed by a C-terminal tail that forms a homodimeric coiled-coil that ends with a short highly-acidic unstructured region that binds GM130. **(B)** Structural model of a p115 dimer showing the head domain and predictions for the coiled-coil tail which is shown in three parts with gaps at the regions predicted to be unstructured and hence likely to be flexible. **(C)** Immunoblot showing expression of GFP-tagged forms of p115 or its separate domains in transfected HEK293T cells, with tubulin as a loading control. **(D)** Confocal micrographs of cells expressing the GFP-tagged forms of p115 shown in (C) and labelled for the ER exit site marker Sec31 and the Golgi marker ZFPL1. Scale bars represent 10 μm. **(E)** Volcano plot comparing enrichment of proteins co-precipitating with p115-GFP versus GFP from HEK293T cells. Values are means of biological triplicates. Highly enriched proteins involved in membrane traffic are indicated in red and named. The other highly enriched proteins are the centriolar components AKAP9, CDK5RAP2, PCNT, PCM1, of which AKAP9 (AKAP450) has been reported to be recruited to the Golgi by binding GM130 (Wu et al., 2016). **(F)** Volcano plot comparing protein enrichment of proteins co-precipitating with of p115-tail-GFP versus p115-head-GFP from HEK293T cells. Values are means of biological triplicates. The other highly enriched proteins with the p115 tail are the pericentriolar proteins noted in (E).

Despite its clear importance in the early secretory pathway the precise role and mechanism of action of p115 are not fully resolved. In vitro reconstitution of ER to Golgi transport in yeast showed that Uso1p acts to tether vesicles to Golgi membranes before the SNAREs act to drive fusion (Barlowe, 1997; Cao, 1998). This step also requires the Rab1 ortholog, Ypt1, and in both yeast and mammals this GTPase binds directly to p115 to promote its association with Golgi membranes, with Rab1 being proposed to bind to either the head domain or the coiled-coil region (Allan et al., 2000; An et al., 2009; Beard et al., 2005; Brandon et al., 2006). It has also been reported that the coiled region of p115 interacts with SNAREs to promote their assembly, although a recent study in fungi suggested that the SNAREs interact with the head domain (Bravo-Plaza et al., 2023; Shorter et al., 2002). To further illuminate the role of p115 we have sought new interaction partners using a proteomic approach and find an unexpected interaction with the Sec16A component of ER exit sites.

## RESULTS AND DISCUSSION

### Identification of interaction partners of p115

To investigate the biological role of p115 we sought interaction partners by immunoprecipitation of GFP-tagged forms of the protein. Full-length p115 was expressed with GFP at its C-terminus, along with two truncations each containing one of the two distinct parts of the protein, the head domain and the C-terminal coiled-coil. These three fusion proteins accumulated to similar levels and were, as expected, at the Golgi with lower levels in the cytoplasm, indicating that they were correctly folded (Fig. 1C,D). Full-length p115-GFP was precipitated, with GFP alone used as a negative control, and associated proteins were identified with mass-spectrometry. Comparison of p115-GFP to the GFP negative control revealed a strong enrichment for two known interactions partners: the Golgi coiled-coil protein GM130 which binds directly to the C-terminus of p115, along with GRASP65 which is known to bind directly to GM130 (Fig. 1E).

In addition to these known interactions, several proteins not previously associated with p115 were strongly enriched in the precipitations including two components of membrane traffic, Sec16A and Sec13. Sec16A is a large protein that organises ER exit sites and promotes COPII vesicle budding by interaction with components of the Sar1:Sec23:Sec24 inner layer of the coat, and Sec13 is a beta-propellor protein that forms a stable complex with Sec16A (Watson et al., 2006; Whittle and Schwartz, 2010). Sec13 also forms a complex with the COPII coat subunit Sec31 to form the outer layer of coat, but the latter protein was not enriched in the precipitates (Downes and Zanetti, 2025). To further investigate these interactions with p115 we compared the proteins that precipitated with the two halves of the protein and found that GM130 and its interactors precipitated with the C-terminal tail as expected whilst Sec16A and Sec13 were the most strongly enriched interactors with the helical head domain (Fig. 1F).

### Mapping the region of Sec16A required for interaction with p115

Sec16A is predicted to be mostly unstructured with a central conserved domain that forms a helical solenoid resembling part of a COPII subunit (Fig. 2A). This region of the protein homodimerises, and each monomer in the complex binds to the beta-propellor protein Sec13, a protein present as a subunit in several other complexes (Dokudovskaya et al., 2011; Whittle and Schwartz, 2010). The C-terminal unstructured region binds to the Sec23 subunit of COPII and its associated Sar1 GTPase and has been reported to accelerate COPII coat formation (Kung et al., 2012; Yorimitsu and Sato, 2012). To determine where p115 binds to Sec16A we expressed in cells a fusion of p115 to the C-terminal mitochondrial targeting signal from monoamine oxidase (MAO). Sec16A is normally located to ER exit sites which are scattered throughout the cytoplasm with an accumulation next to the Golgi apparatus (Watson et al., 2006). In the presence of mitochondrial p115, both endogenous and exogenous Sec16A relocate to the surface of mitochondria (Fig. 2B,C). The Golgi apparatus is fragmented by the presence of this mitochondrial p115, presumably due to the depletion of endogenous Sec16A and other p115 interaction proteins. Nonetheless, this in vivo relocation provides a means to map the part of Sec16A that interacts with p115, and series of truncations of Sec16A were expressed along with a mitochondrial form of the p115 head domain (Fig. 2A and D). Residues 1-1101 of Sec16A are predicted to be primarily unstructured but were relocated by the mitochondrial p115 (Fig. 2D). In contrast, the rest of the protein which contains the central conserved region and the Sec23 binding site was not relocated when expressed either in its entirety or as fragments containing either the CCD or the Sec23 binding site (1101-1890 or 1890-2357; Fig. 2D). Truncation of the N-terminal region showed that residues 511-1100 were still relocated by ectopic p115. To further map the region of binding, region 511-1100 was split into thirds and each fragment expressed as a GFP fusion and then precipitated from cells. Immunoblotting showed that the central region, residues 650-776 was found to be sufficient for co-precipitating p115 (Fig. 2E). Taken together, these results show that the head domain of p115 can interact with Sec16A via a 126-residue region in the latter’s N-terminal unstructured region.

**Figure 2.**
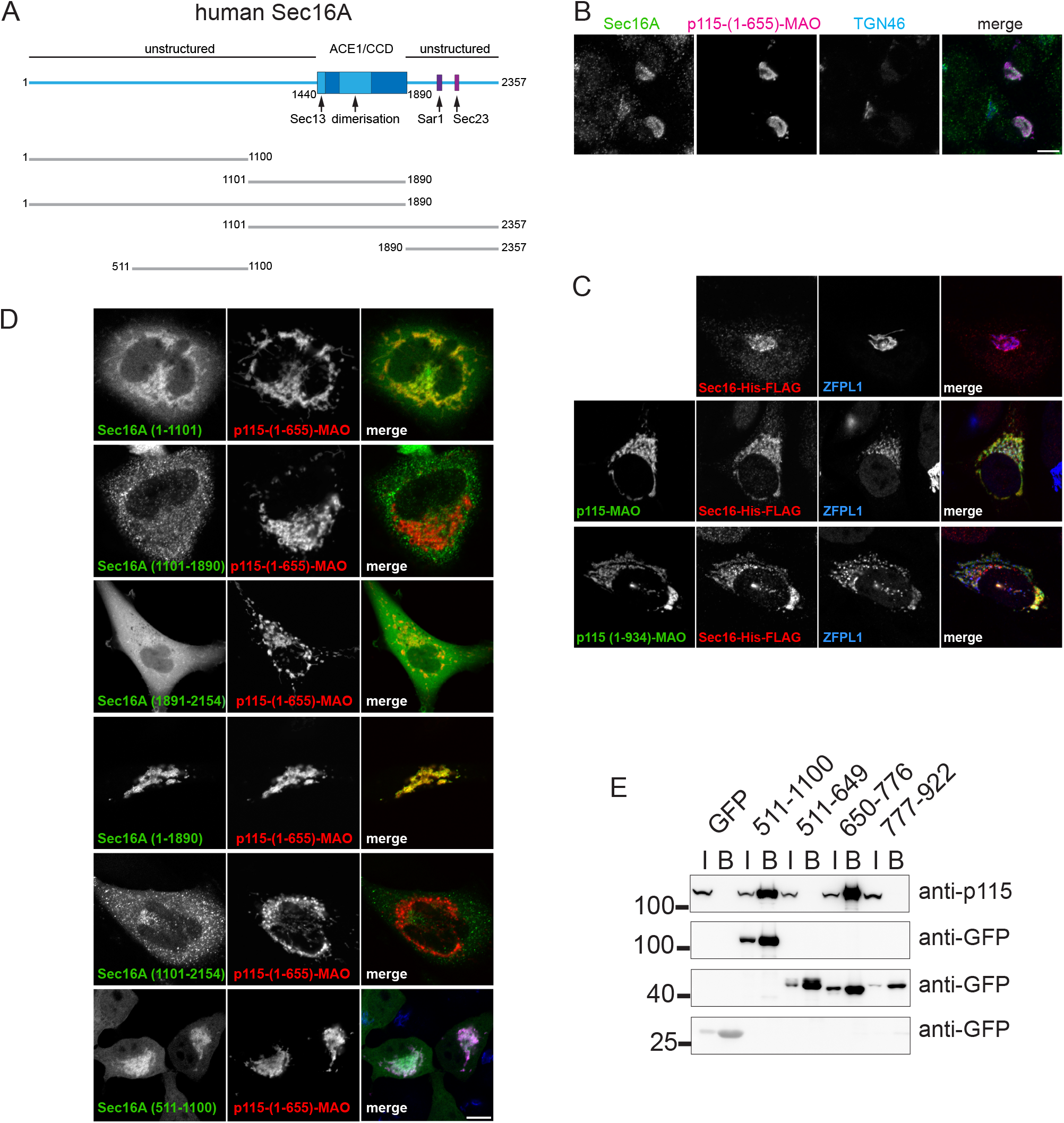
p115 interacts with Sec16A via part of the N-terminal unstructured region. **(A)** A cartoon schematic of human Sec16A showing the truncations used to map the p115 binding site. Most of the protein is predicted to be unstructured apart from the central conserved domain (CCD, also referred to as the ancestral coatomer element 1 (ACE1)), which comprises a β solenoid that forms a homodimer, along with a short region of sheet that binds the β-propellor protein Sec13 (Bhattacharyya and Glick, 2007; Whittle and Schwartz, 2010). The C-terminal unstructured region binds to Sec23 and its associated Sar1 GTPase (Bhattacharyya and Glick, 2007; Yorimitsu and Sato, 2012). **(B)** Confocal micrographs showing the capture of endogenous Sec16A by p115-HA-MAO on mitochondria in HeLa cells with TGN46 acting as a Golgi marker. **(C)** Confocal micrographs of HeLa cells expressing p115-HA-MAO and Sec16A-His6-FLAG, and labelled for the HA and His6 tags along with the Golgi marker ZFPL1. **(D)** Confocal micrographs of HeLa cells expressing p115(1-655)-HA-MAO and the indicated fragments of Sec16A fused to GFP and labelled for the HA tag. **(E)** Immunoblots showing interaction of endogenous p115 with the indicated GFP-tagged fragments of Sec16A expressed in HEK293T cells and then precipitated (I: input, B: bound).

### Structural prediction reveals the basis of a direct interaction between p115 and Sec16A

To obtain more insight into the interaction between p115 and the N-terminal region of Sec16A we used the AlphaFold 3 structural prediction algorithm (Abramson et al., 2024). When the whole N-terminal region of Sec16A (residues 1-1100) was tested with the head domain of p115, two short regions of Sec16A gave a very confident prediction for binding (Fig. 3A). These two regions both lie within the 650-776 binding region mapped as described above (residues 718-726 and 738-746). The two regions are well conserved in vertebrates but do not appear to be conserved in invertebrates (Fig. 3B). In the predicted complex with p115 these regions are part of an unstructured region which binds in a groove on the surface of the head domain, with this groove being on the opposite side to the attachment point for the coiled coil (Fig. 3C). To test this predicted structure, we examined the effect of mutating residues in Sec16A that appear to be involved in the interface with p115. When a short stretch of residues in either the first region (site 1) or the second region (site 2) was mutated to alanine then region 650-776 lost the ability to co-precipitate endogenous p115 when expressed as a GFP fusion in transfected cells (Fig. 3D). When the region 650-776 of Sec16A was expressed on the surface of mitochondria endogenous p115 relocated to the same organelle, and again this activity was lost with either the site 1 or site 2 mutations (Fig. 3E). These results indicate that a conserved bi-partite motif in the N-terminal unstructured region of Sec16A can bind directly into a groove in the head domain of p115.

**Figure 3.**
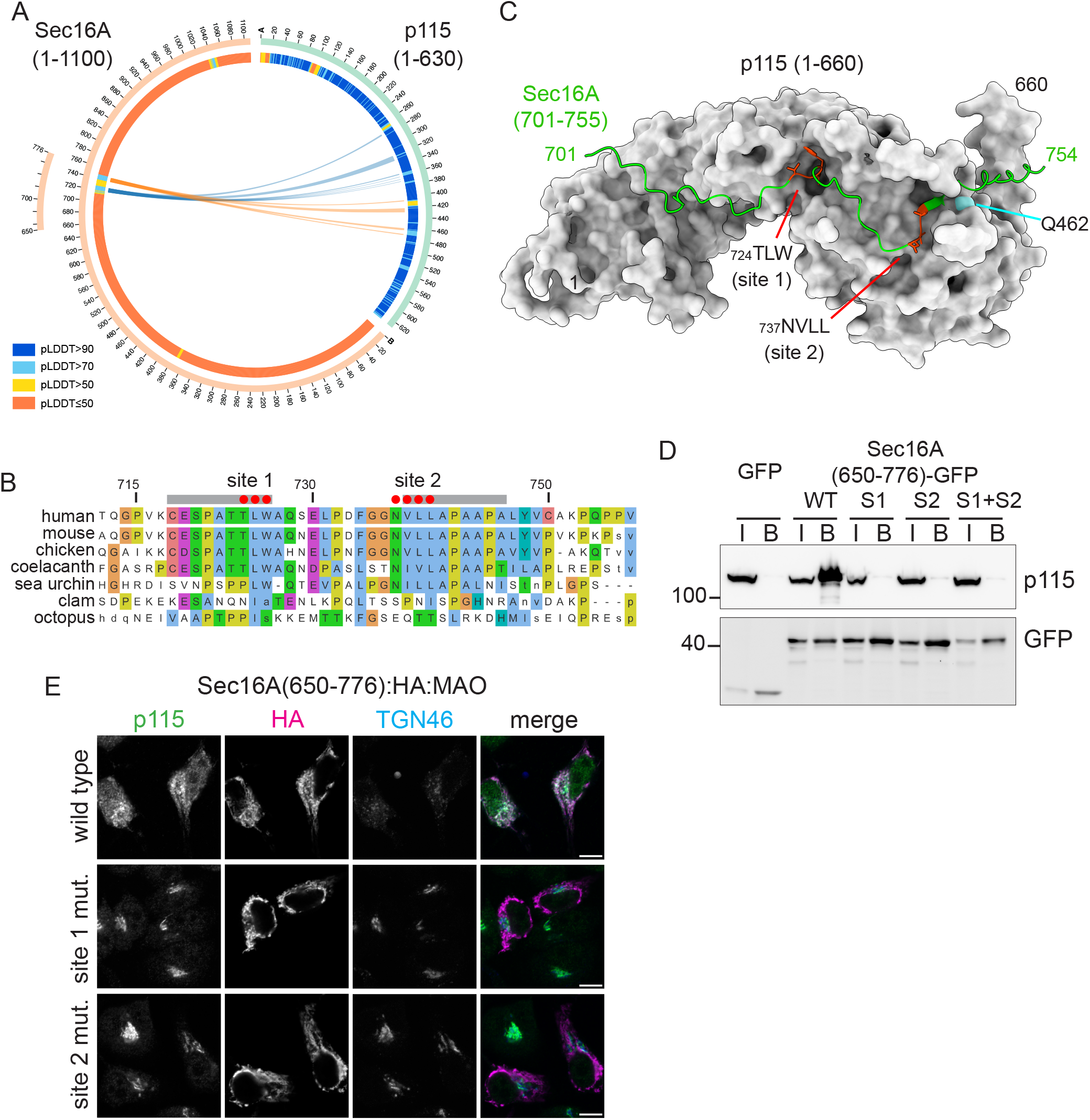
Prediction of a complex between Sec16A and the head domain of p115. **(A)** AlphaBridge visualisation of the output from an AlphaFold 3 prediction of a complex between the N-terminal region of Sec16A (1-1100) and the head domain of p115 (1-630). Shown in orange and blue are the two high confidence interactions identified by AlphaBridge, both of which have a predicted interaction Confidence Score (piCS) of 0.91 (Álvarez-Salmoral et al., 2025). The pLDDT score (≤50) indicates that most of this region of Sec16A is predicted to be unstructured. **(B)** Proviz alignment of human Sec16A (712-757) showing conservation in the indicated vertebrates and invertebrates. The two regions strongly predicted to interact with p115 are indicated in grey along with the residues that were selected for mutation (red). These regions are well conserved in vertebrates but not in invertebrates. **(C)** AlphaFold 3 prediction of the complex formed between residues 701-754 of Sec16A and the head domain of p115. The residues of Sec16A lie along a groove in the surface of p115, with two regions predicted to interact with very high confidence. Shown in red are short stretches of residues in the middle of each region that were selected for mutation to alanine. **(D)** Immunoblot of anti-GFP precipitations from cells expressing the indicated forms of Sec16A(650-776), or GFP alone as a negative control, and then probed for p115 and GFP. **(E)** Confocal micrographs of HeLa cells expressing the indicated mitochondrial forms of Sec16A(650-776), and probed for endogenous p115, the Golgi marker TGN46 and the HA tag in the mitochondrial chimera. Scale bars represent 10 μm.

### Sec16A binding is required for full activity of p115

To test the role of Sec16A binding on the activity of p115 we tested the effect of perturbing the groove in which Sec16A is predicted to bind. A glutamine at residue 462 lies against Sec16A in this binding grove (Fig. 3C). Mutation of this residue to aspartate resulted in loss of binding to Sec16A (Fig. 4A,B). When expressed in cells, this mutant form still localised to the Golgi and when precipitated from cells it was associated with similar levels of GM130 (GOLGA2) to that found with wild-type p115 indicating that the mutant form was not grossly misfolded (Fig. 4C,D). To test its activity in vivo we used an assay for anterograde traffic that is based on releasing a GFP-tagged membrane protein from the ER which then traffics to the cell surface where it can be detected with a fluorescent antibody and hence its traffic quantified by flow cytometry (Boncompain et al., 2012; Pereira et al., 2023) (Fig. 4E). The USO1 gene encoding p115 is essential for cell viability, but it can be transiently disrupted by delivery of CRISPR-Cas9 guides with lentivirus, and as expected this treatment blocked traffic of the reporter protein to the surface (Fig. 4F). To test the functional activity of p115, Halo-tagged forms expressed from a guide-resistant cDNA were transfected back into the CRISPR-Cas9 treated cells with the transfected cells in the population being identified with flow cytometry. Wild type p115 gave good rescue of the p115 disruption but this rescuing activity was reduced by the Q462D mutation (Fig. 4G). The reduction in rescuing activity was similar to that observed when the GM130-binding site on p115 is removed, an interaction made by p115 that is very well established but known to be not absolutely essential for its activity (Fig. 4F,G). Taken together, these results show that the Sec16A binding site in p115 is functionally important for optimally efficient membrane trafficking.

**Figure 4.**
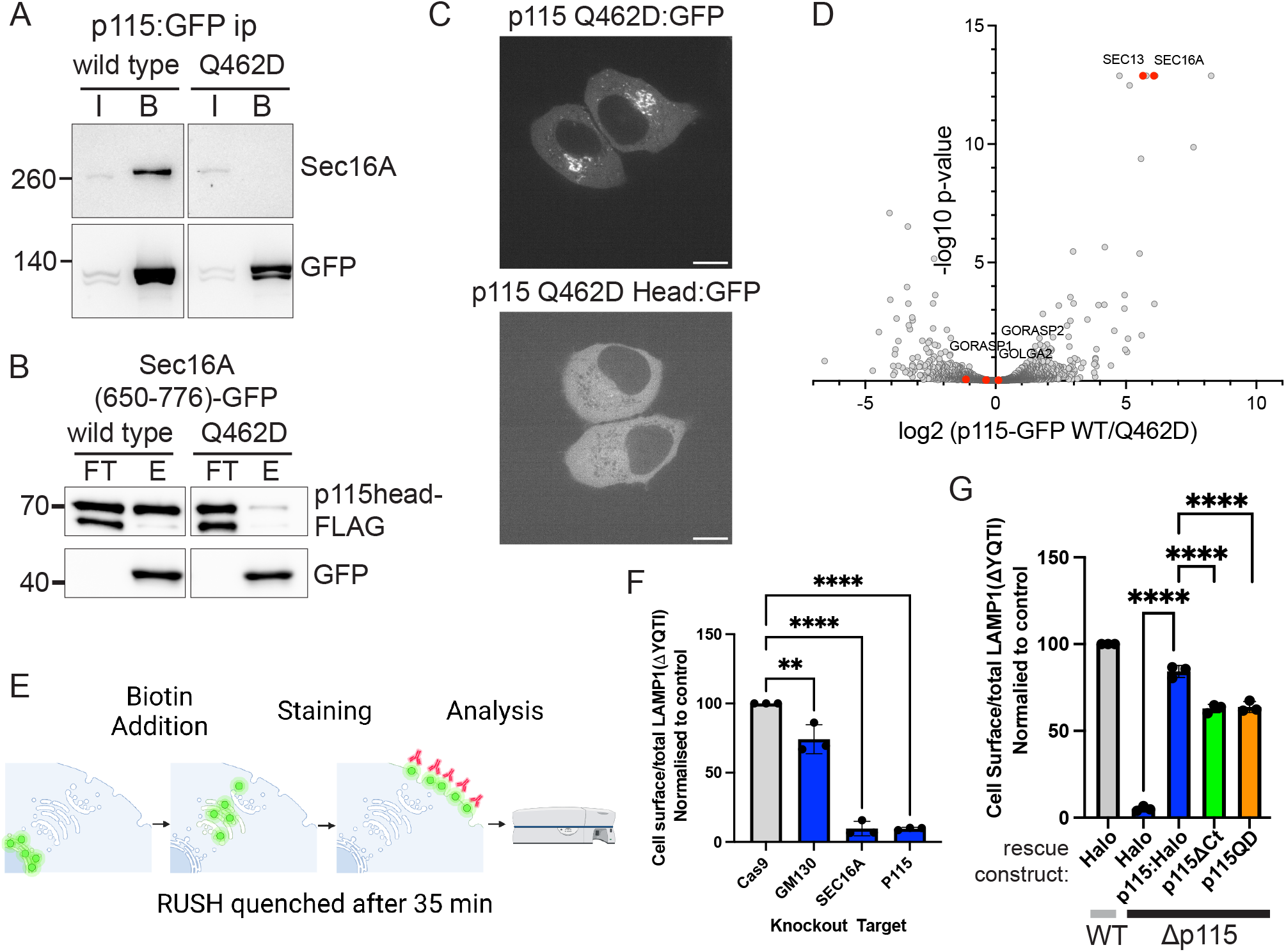
Mutation of the Sec16A binding site in p115 reduces its activity. **(A)** Immunoblot showing binding between endogenous Sec16A and either wild-type full-length p115-GFP or the same with the Q662D mutant expressed in HEK293T cells and precipitated with anti-GFP (I, input; B, bound). **(B)** Immunoblot of a direct binding assay performed using flag-tagged p115 head domain (1-655) binding to Sec16A(650-776)-GFP on GFP-trap beads (FT, flow-through/unbound; E, eluate/bound). **(C)** Confocal micrographs of live HeLa cells expressing full-length p115-GFP Q462D fused to GFP, or the head domain (1-655) with the same mutation and tag. Scale bars represent 10 μm. The head domain shows clearer Golgi labelling by live cell imaging than by immunofluorescence (Fig. 1D), indicating that the targeting is partially lost during fixation. **(D)** Volcano plot comparing enrichment of proteins co-precipitating co-precipitating with wild-type p115-GFP vs to Q462D p115-GFP following transfection into HEK293T cells. Values are means of biological triplicates. Proteins found enriched with wild-type p115 are indicated, showing that the interaction with Sec16A and Sec13 are the ones that change with the mutant whilst GOLGA2 and its binding partners are unaffected. **(E)** Schematic of the flow cytometry RUSH assay. Green circles represent LAMP1ΔYQTI-GFP cargo, red antibodies represent mCherry-tagged anti GFP nanobodies. LAMP1 is held in the ER by an SA-KDEL hook which is released upon biotin addition allowing LAMP1 to be transported to the cell surface where it can be quantified by antibody binding and flow cytometry. **(F)** Bar graph showing relative effects of GM130, p115 and Sec16A loss on anterograde trafficking (n=3), error bars indicate SD. Statistical comparisons were made by one-way ANOVA testing, **** = p<0.0001, ** =p<0.01 (grey - untreated Cas9 cells as control, blue – the same cells infected with lentivirus expressing CRISPR guides). **(G)** Bar graph showing the degree of rescue of traffic in cells lacking p115 by either wild-type or mutant p115. The first column represents a Cas9 cell line transfected with Halo (WT), while all other columns represent the same cells treated with lentivirus expressing a CRISPR guide for p115 (Δp115) and transfected with Halo as a control or the indicated p115-Halo constructs (n=3).

### Conclusions

In this paper we show that the head domain of p115 can bind directly to a motif in the unstructured N-terminal region of Sec16A. This interaction does not appear to be responsible for the essential role of p115 in secretion in cultured cells. p115’s essential role seems more likely to be that of promoting the assembly of SNARE complexes on the early Golgi, potentially directed by Rab1, although the details of this remain to be resolved (Allan et al., 2000; An et al., 2009; Beard et al., 2005; Brandon et al., 2006). Indeed, the pattern of conservation of the sequence recognised by p115 suggests that this interaction arose early in vertebrate evolution as an extra function of p115 in addition to its function in vesicle tethering (Fig. 3B). Invertebrates and fungi only have a single Sec16 gene, but early in vertebrate evolution this was duplicated to form Sec16A and Sec16B (Bhattacharyya and Glick, 2007; Budnik et al., 2011). Sec16B does not have the p115 binding motif described here, but its in vivo function is unclear as it is only expressed at low levels in a limited range of cell types and it is not essential for viability or fertility in mice, in contract to Sec16A which is essential at a cellular level (Groza et al., 2023).

We speculate that p115 binding to Sec16A could provide a link between ER exit sites and the ERGIC/cis Golgi. The C-terminus of p115 is known to bind the N-terminus of the early Golgi golgin GM130, but the purpose of this is unclear as this part of p115 is not essential for its role in secretion (Nelson et al., 1998; Puthenveedu and Linstedt, 2004). The long regions of coiled-coil in both proteins would allow the head domain of p115 to extend at least 200 nm from the ERGIC/cis Golgi membrane on which GM130 is anchored at its C-terminus. Binding Sec16A could potentially enable a population of p115 molecules to hold the ER exit sites and early Golgi in close proximity, or alternatively it could help attach ER-derived vesicles to the tethers on their destination membrane whilst they are still in the process of budding. Further work will be required to resolve these possibilities, but it seems likely that investigating this, and other interactions made by the machinery at ER exit sites, will reveal the mechanisms by which this fundamentally important region of the cell is organised.

## MATERIALS AND METHODS

### Cell culture and transfection

HeLa, HEK293T (ATCC) and LAMP1ΔYQTI RUSH HeLa cells (Pereira et al., 2023) were grown in Dulbecco’s Modified Eagle Medium (DMEM) supplemented with 10% fetal bovine serum (FBS) and penicillin-streptomycin at 37°C and 5% CO2. The cells were regularly checked for mycoplasma (MycoAlert, Lonza Biosciences). For small scale plasmid transfections, cells were split into 6-well plates at a 1:6 dilution. After 24 h, cells were transfected with 1 µg of DNA, 4 µl of 1 mg/ml polyethylenimine (PEI) and 100 µl of Opti-MEM (Thermo Fischer). Either 12 or 24 h later cells were plated onto slides for immunofluorescence or into chambered coverslips for live cell imaging. For T75 flasks, 10 µg of DNA, 30 µl of PEI, and 800 µl of Opti-MEM were used, and for T175 flasks, 25 µg of DNA, 75 µl of PEI and 2 ml of Opti-MEM.

### Fluorescence microscopy

For immunofluorescence, cells were plated onto multi-well PTFE-coated slides (Hedley-Essex), 12-24 h after transient transfection and placed in a humidified chamber inside a 37°C 5% CO_2_ incubator for 12-24 hours. Slide wells were washed with PBS, fixed for 20 min in 4% PFA in PBS, rinsed three times with PBS, and permeabilised for 10 min in 0.5% Triton X-100 in PBS. After washing twice in PBS, cells were treated with blocking buffer (PBS, 4% FBS and 1% Tween-20) and incubated in blocking buffer containing primary antibodies for 1 h. The wells were washed thrice with PBS and incubated with secondary antibodies in blocking buffer for 1 hour in the dark. After washing thrice with PBS, cells were mounted under a coverslip in VECTASHIELD with DAPI (Vectorlabs) and sealed with nail varnish. Images were obtained with on a Zeiss 780 confocal and processed using Fiji. Primary antibodies were against Sec16A (ab70722, Abcam); Sec31A (612350, BD Biosciences); ZFPL1 (HPA014909, Sigma-Aldrich); p115/USO1 (ab184014, Abcam); HA tag (ROAHAHA, Sigma-Aldrich) and the His tag (ab18184, Abcam), and were detected with species-specific AlexaFluor-labelled secondary antibodies. For live cell imaging, cells were seeded into µ-slide chambered coverslips with ibiTreat coating (Ibidi), in 300 µl of media 24 hours prior to imaging on a Nikon iSIM microscope with the stage kept at 37°C and 5% CO_2_.

### Precipitation of GFP-tagged proteins

One T175 flask of sub-confluent HEK293T cells was transfected with a plasmid expressing the GFP-tagged protein and after two days the cells were detached and pelleted by centrifugation. The pellet was resuspended in 500 µl lysis buffer (10 mM Tris-HCl pH 7.5, 150 mM NaCl, 0.5 mM EDTA, 0.5% TX-100, cOmplete Protease Inhibitors (Roche)) and incubated for 30 min at 4°C. The lysate was cleared at 16,000 x g for 15 min, 25 µl was saved as an input control, and the rest incubated at 4°C for one hour with 50 µl GFP-trap agarose bead slurry (Chromotek) that had been washed twice with 500 µl lysis buffer. At the end of incubation, 25 µl of supernatant was saved (unbound fraction), and the beads washed three times in wash buffer (10 mM Tris pH7.5, 300 mM NaCl, 0.5 mM EDTA, cOmplete Protease Inhibitors (Roche)). 60 µl of 2x sample buffer was added to the washed beads, and after heating at 100 °C for 5 min the beads were pelleted and the supernatant was transferred to a fresh tube and BME was added to a concentration of 10%. For mass spectrometry analysis, 30 µl were separated on a 12-well 4-20% tris-glycine gel (ThermoFisher), stained with InstantBlue (Sigma-Aldrich) for 1h and each lane cut into 8 equal slices for trypsinisation and protein identification by label-free mass spectrometry. Analysis of samples by immunoblotting used primary antibodies against Sec16A (HPA005684, Sigma-Aldrich); p115 (ab184014, Abcam); tubulin (YL1/2); FLAG tag (F3165, Sigma-Aldrich); HA tag (3F10, Roche); His tag (18148, Abcam); and GFP (SAB4301138, Sigma-Aldrich).

### Protein binding assays

Two populations of HEK293T cells were used: one transfected with a plasmid expressing a GFP-tagged protein and the second expressing a FLAG-tagged protein. After 48 h, the cells were detached and lysed in 1 ml of lysis buffer as described above for GFP-tagged proteins. Lysates were incubated at 4°C for 1 hour with either 100 µl of Anti-FLAG M2 Affinity Gel (Merck) used per T175 flask for FLAG-tagged protein or 40 µl of GFP-Trap beads for GFP-tagged proteins, with both sets washed twice in 500 µl lysis buffer prior to addition of lysates. The GFP-Trap beads underwent a two-step harsh wash (2x Wash 1: 1% SDS in water; then 3x Wash 2: 50 mM Tris pH 7.4, 1 M NaCl, 0.5 mM EDTA, 1% Triton X-100). Anti-FLAG beads were washed three times with lysis buffer, with the second wash supplemented with 1 M NaCl, and then incubated with 500 µl elution buffer (25 mM Tris-Hcl pH 7.4, 250 mM NaCl, 1mM EDTA, 100 µg/ml 3xFLAG peptide) for 15 min at room temperature. The eluted FLAG protein was incubated with the GFP-tagged protein on GFP-trap beads for 4°C for 1h. The beads were then washed three times with original wash buffer (10 mM Tris pH7.5, 300 mM NaCl, 0.5 mM EDTA, cOmplete Protease Inhibitors (Roche)) and eluted as for the GFP precipitation.

### RUSH secretion assay following lentiviral-based gene knockout

Oligonucleotide sequences corresponding to suitable guides were cloned into the pLVK lentiviral vector (Takara Bio) between two BbsI sites. 1.25×10^6^ Lenti-XTM 293T cells (Sigma-Aldrich) were seeded per well of a 6-well plate coated with matrigel. 240 µl Opti-MEM, 1 µg of each guide plasmid, 1.6 µg of Gag/Pol plasmid and 0.4 µg of VSVG plasmid was combined with 240 µl of Opti-MEM containing 10 µl of Lipofectamine 2000, incubated for 20 min and then added the cells. Cell supernatants were collected after 4 days, passed through a 0.45 µm filter and stored at -80 °C. For infection, 50 µl of 5×10^4^ suspension of Cas9-expressing LAMP1ΔYQTI HeLa cells were transduced with 200 µl of lentivirus suspension and centrifuged at 700 x g at 37°C for 1 h in a 48 well plate. After 24 hours, cells were transferred from the 48 well plate to a 6 well plate. For rescue experiments, two 48-well plate wells were used for each rescue construct, and the cells were seeded into a 10 cm plate instead of a 6-well plate. The p115 rescue constructs were engineered to contain silent mutations to prevent recognition by the p115 CRISPR guides. Six days post-infection, cells were trypsinised, placed in DMEM, washed three times with 3 ml Opti-MEM by centrifugation, and resuspended in 200 µl Opti-MEM. 10^6^ cells in 200 µl of opti-MEM were added to 0.5 µg of DNA and electroporated with the Gene Pulser Xcell electroporator (BioRad) and the default HeLa electroporation protocol. Cells resuspended in 2 ml DMEM were transferred to 6 well plates, 1 ml of DMEM containing Halo-646 dye (Janelia) was added to a final concentration of 20 nM and the cells incubated overnight.

For the RUSH assay, one week after infection, cells were trypsinised, suspended in 5 ml DMEM, and centrifuged for 5 min at 500 xg. The cells were resuspended in 1 ml CO_2_ free media (DMEM + 25mM HEPES pH7.4), split equally between two 1.5 ml tubes and 0.5 ml of CO_2_-free media added to one (t=0). All tubes were placed in a 37°C hot block, 1 ml CO_2_-free media with1 mM biotin added to the tubes apart from t=0, and RUSH allowed to progress for 35 min. Cells were centrifuged, resuspended in 100 µl of PBS containing 10 µg/ml mCherry anti-GFP nanobody for 1h, then washed twice in 0.5 ml PBS, resuspended in 0.4 ml PBS and were passed through a 35 µm cell strainer for flow cytometry (LSRFortessa). Data from three repeats was processed using FlowJo and ANOVA testing used to compare the differences between cell populations (GraphPad Prism 9).

### Computational tools and methods

For mass spectrometric data the DEP R package (v1.30.0) was used to analyse the LFQ intensities of each protein identified across the samples (Zhang et al., 2018). The data was filtered to exclude contaminants and reverse sequences. Any duplicated proteins were removed. Each sample was grouped with the two other repeats for each condition. A filter was applied with the rule that a protein had to be present across all three repeats of a condition to be included. The data was normalised and missing values were added by imputation. We opted to use a left-shifted Gaussian distribution with a down-shift of 1.8 standard deviations and width of 0.3 standard deviations relative to the parent population. This generates a small and more tightly spaced distribution of normalised LFQ intensities. Missing values are then randomly picked from the distribution to allow calculation of the enrichment values and these compared for each protein between samples to derive a p-value. Protein alignments were with ProViz, and structural prediction was with AlphaFold3 followed by analysis in AlphaBridge and visualisation in ChimeraX (Abramson et al., 2024; Álvarez-Salmoral et al., 2025; Jehl et al., 2016; Pettersen et al., 2020).

## Acknowledgements

Mass spectrometry analysis was performed at the Biological Mass Spectrometry and Proteomics Facility of the MRC LMB. Funding was provided to SM by the Medical Research Council, as part of United Kingdom Research and Innovation (also known as UK Research and Innovation) file reference number MC_U105178783; and to DG by Biotechnology and Biological Sciences Research Council responsive mode grants BB/W005905/1 and UKRI715.

